# Prioritization of SARS-CoV-2 epitopes using a pan-HLA and global population inference approach

**DOI:** 10.1101/2020.03.30.016931

**Authors:** Katie M. Campbell, Gabriela Steiner, Daniel K. Wells, Antoni Ribas, Anusha Kalbasi

## Abstract

SARS-CoV-2 T cell response assessment and vaccine development may benefit from an approach that considers the global landscape of the human leukocyte antigen (HLA) proteins. We predicted the binding affinity between 9-mer and 15-mer peptides from the SARS-CoV-2 peptidome for 9,360 class I and 8,445 class II HLA alleles, respectively. We identified 368,145 unique combinations of peptide-HLA complexes (pMHCs) with a predicted binding affinity less than 500nM, and observed significant overlap between class I and II predicted pMHCs. Using simulated populations derived from worldwide HLA frequency data, we identified sets of epitopes predicted in at least 90% of the population in 57 countries. We also developed a method to prioritize pMHCs for specific populations. Collectively, this public dataset and accessible user interface (Shiny app: https://rstudio-connect.parkerici.org/content/13/) can be used to explore the SARS-CoV-2 epitope landscape in the context of diverse HLA types across global populations.

## Introduction

Infection with SARS-CoV-2 can result in a spectrum of clinical phenotypes encompassed by COVID-19, from asymptomatic illness to a potentially lethal disease with hallmarks of acute respiratory distress syndrome (ARDS) (Yang et al., 2020). While some clinical demographics have been associated with a more severe disease course (Guan et al., 2020), the heterogeneity of clinical outcomes is otherwise poorly understood.

The range of clinical outcomes may at least in part be related to patient-specific antiviral T cell responses. T cells are crucial for viral clearance and development of immunologic memory (Wherry and Ahmed, 2004) and are plausible contributors to immunopathology following viral infection (Channappanavar and Perlman, 2017). Both SARS-CoV-2 reactive CD4 and CD8 T cells have been detected in patients with COVID-19 (Chour et al., 2020; Grifoni et al., 2020a; Weiskopf et al., 2020a), though early studies suggest the relationship between T cell responses and the severity of COVID-19 is complex (Mathew et al., 2020).

The heterogeneity in T cell responses to SARS-CoV-2 may be related to recognition of viral antigens in the context of class I and II human leukocyte antigen (HLA) proteins (Chour et al., 2020). Indeed, genetic susceptibilities to viral infection have been tied to variation in the major histocompatibility complex (MHC) genes that encode HLA proteins (Dutta et al., 2018; Hill, 2001). Meanwhile, functional differences in viral antigen-specific T cell responses in symptomatic and asymptomatic patients may also contribute to the biology of at-risk populations (Mathew et al., 2020; Weiskopf et al., 2020a).

Further understanding of virus-specific T cell responses may aid in designing and monitoring the impact of preventative SARS-CoV-2 T cell vaccines. In contrast to SARS-CoV-2 vaccines focused on generating antibody responses against the surface spike glycoprotein that facilitates viral entry into the cell (Thanh Le et al., 2020) T cell vaccines have the capacity to generate immune responses against the entire viral proteome (Gilbert, 2012). In fact, non-spike T cell responses may be associated with less severe COVID19 (Peng et al., 2020).

To evaluate patient-specific T cell responses, recent studies have used large pools of SARS-CoV-2 epitopes based on homology with SARS-CoV, or based on prediction of MHC class I- and class II-binding peptides across common HLA alleles in order to capture a broad population (Grifoni et al., 2020b; Smith et al., 2020). To facilitate a more comprehensive evaluation of anti-viral and vaccine-induced T cell responses, and to support region-specific and global vaccine design strategies, we generated a resource database with a corresponding user-friendly interface to facilitate exploration of predicted MHC-binding peptides across 9,360 and 8,445 class I and II HLA alleles, to account for the genetic diversity in the MHC gene complex across global populations.

## Results

### In silico predictions of SARS-CoV-2 antigens

We deployed binding predictions across the SARS-CoV-2 proteome (**Figure 1**) for 9,360 class I HLA alleles (2,987 HLA-A; 3,707 HLA-B; 2,666 HLA-C; 9-mers) and 8,445 class II HLA alleles (15-mers). The predicted binding affinity (in nanomolar [nM]) between peptides and HLA proteins (pMHCs) were summarized by the median predicted binding affinity across all algorithms (Median Score). The Median Score values were filtered to those less than 500nM, a common filter used in peptide binding predictions for the purpose of identifying T cell epitopes (Rajasagi et al., 2014; Sidney et al., 1999). There were 368,145 unique combinations of peptides and HLA alleles (pMHCs) with a predicted binding affinity of less than 500nM (**Table S2**), including 1,103 unique 9-mer and 2,547 15-mer peptides and 1,022 MHC class I and 3,481 MHC class II HLA proteins, respectively. Of note, 905 9-mers (82%) were nested within 1,789 15-mers (70%), indicating that a subset of peptides are predicted as both class I and II epitopes.

**Figure 1.**
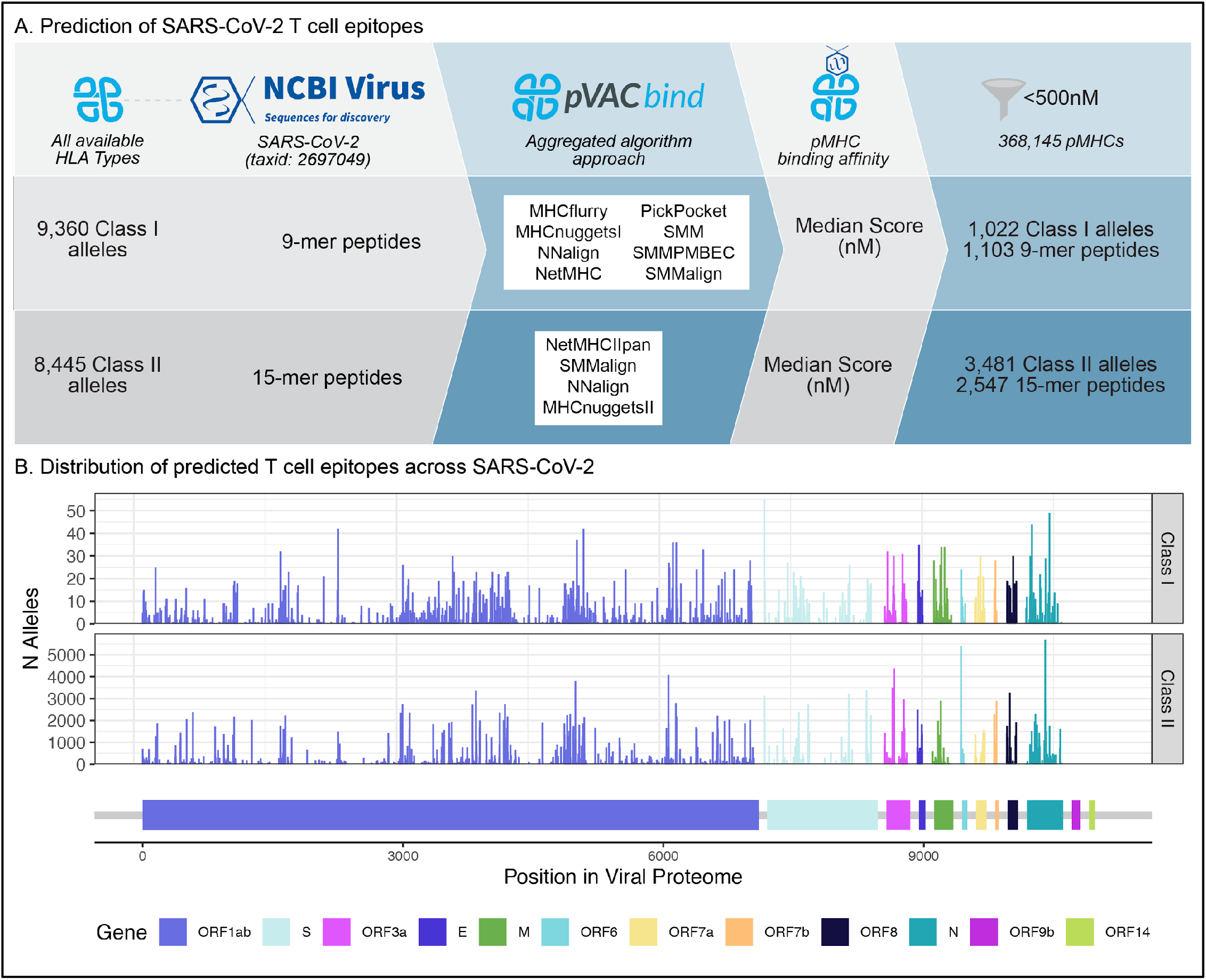
Peptide binding predictions for SARS-CoV-2. A. Overview of the analysis strategy. Class I and II HLA alleles, combined with 9-mer and 15-mer peptides spanning the viral proteome were used as inputs for an aggregated peptide binding prediction approach. A filter of peptides with a median score of 500nM was applied to summarise a set of peptide-MHC complexes (pMHCs) with predicted high binding affinity. B. The distribution of the number of HLA alleles (distinguished by Class I vs II) is shown, according to corresponding peptide, indicated by its starting position within the viral proteome (x-axis). Peptides are colored by their corresponding genes.

In order to better understand the predicted antigenic profile of SARS-CoV-2, we focused on the set of 368,145 pMHCs with predicted binding affinity of less than 500nM for the subsequent analyses. Both class I and class II antigens were predicted across 10 of the SARS-CoV-2 genes (**Figure 1B**), with the most derived from *Orf1ab* (n=690 9-mers; 1,589 15-mers), encoding the Orf1ab polyprotein. The number of peptides from each gene correlated with protein length (R^2^ = 0.997, p=2.10e-11; **Figure S1**).

### Confirmation of predicted SARS-CoV-2 antigens in published datasets

In order to assess the validity of the predictions in our dataset, we compared our predicted antigens to previously reported SARS-CoV-2 or SARS-CoV T cell epitopes. There were 9 nine-mer and 5 fifteen-mer peptides in our dataset that were previously validated experimentally as T cell epitopes and reported in IEDB from SARS (**Table 1**). Since our dataset was restricted to 9-mers and 15-mers, we expanded this search to include any IEDB epitopes that overlapped (i.e. either nested, or in overlapping positions) with our predicted peptides, which resulted in 81 additional epitopes (**Table S1**). Four of these total 95 epitopes were specifically associated with HLA-A*02:01, while HLA restrictions were not reported for the remaining 91 epitopes. Each of the 154 peptides from our dataset overlapping with the 95 epitopes reported in IEDB were each predicted to bind a median of 4 class I HLA proteins (range 1-49) and 35 class II HLA proteins (range 1-5,694), suggesting these experimentally validated epitopes may be relevant in multiple HLA contexts.

**Table 1.**
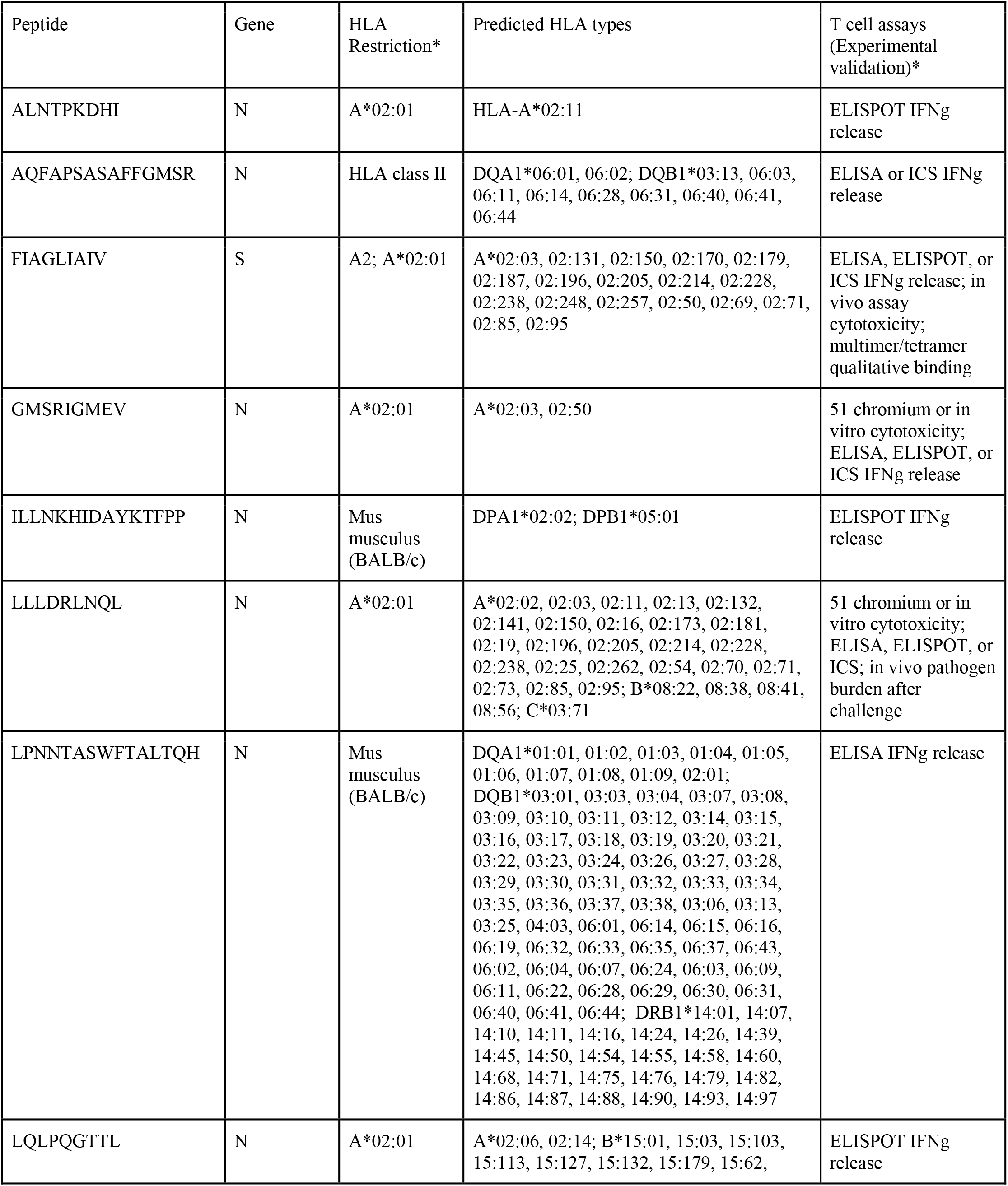
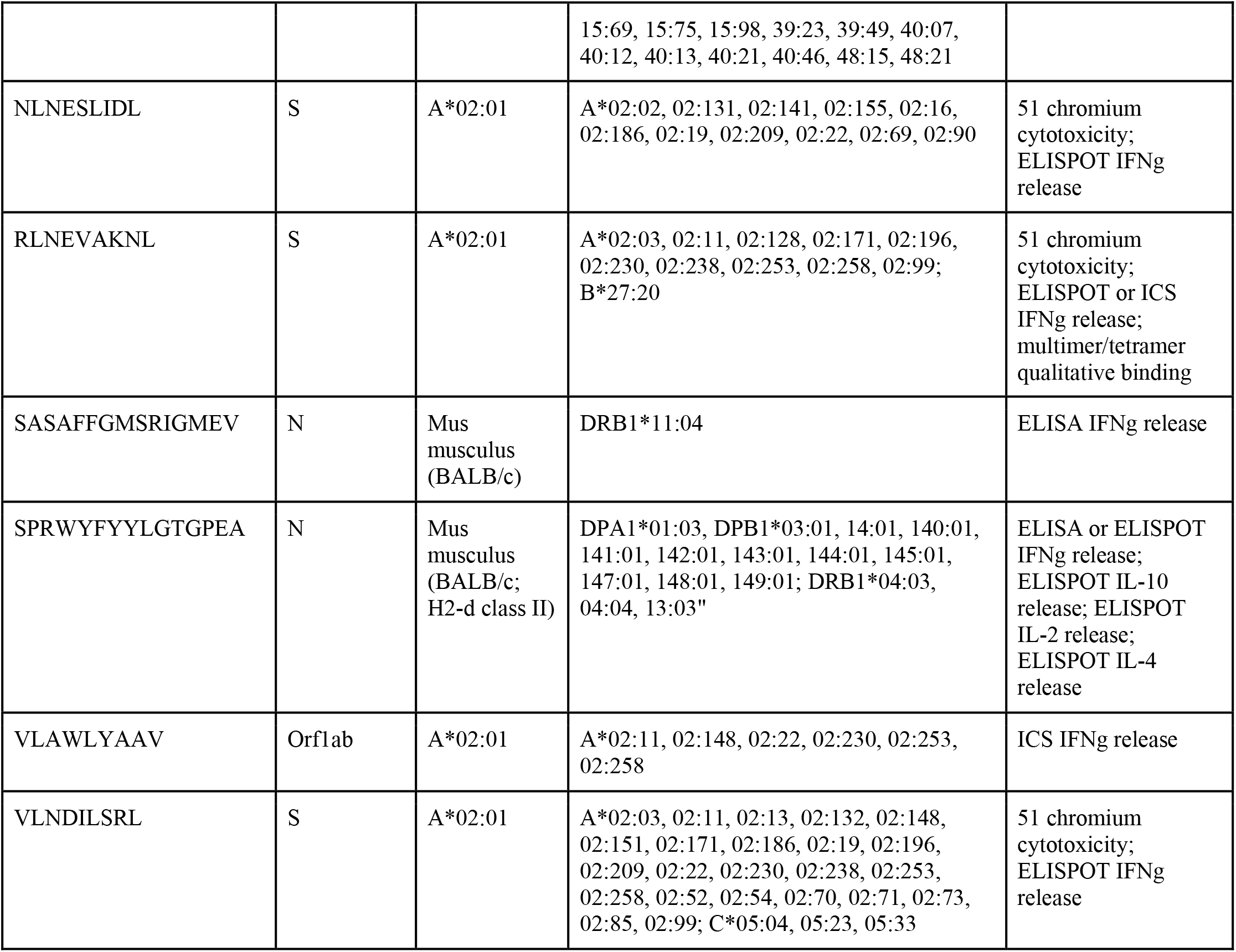
Previously validated T cell epitopes in SARS-CoV from IEDB. *HLA Restriction and Experimental validation are taken from the annotation reported in IEDB. ICS: intracellular cytokine staining.

Grifoni et al. recently used the homology between the SARS-CoV and SARS-CoV-2 proteomes and existing annotated epitopes of SARS-CoV from IEDB to infer T cell epitopes derived from SARS-CoV-2 (Grifoni et al., 2020b). This pool of peptides was assessed in samples derived from COVID19 patients, resulting in the identification of SARS-CoV-2-associated CD4 and CD8 T cell responses in 100% and 70% of convalescent COVID19 patients, respectively (Grifoni et al., 2020a). Our dataset identified 271 nine-mer peptides and 331 fifteen-mer peptides that either overlapped or were nested in 241 CD8 and 628 CD4 T cell epitopes from this study, derived from 9 SARS-CoV-2 genes (**Table S1**). Still, there were 793 nine-mer and 2,139 fifteen-mer peptides in our dataset not included in the megapools experimentally evaluated in this study. Including these additional peptides in experimental validation may increase the sensitivity of detection of T cell responses in patients with SARS-CoV-2.

### Accounting for regional and global relevance of predicted class I pMHCs

To address the regional and global relevance of our predicted class I pMHCs, we aggregated class I HLA frequency data from the Allele Frequency Net Database (AFND) (Gonzalez-Galarza et al., 2020), representing 77 countries from 11 global regions (**Table S2**). Simulated populations (n=100,000 individuals) were created for each individual country, as well as an additional “global” population, constructed by the weighted population frequency of HLA types across countries. Each simulated individual was mapped to their predicted epitope profile by matching their HLA types to their corresponding predicted pMHCs.

Restricting our analysis to HLA alleles with at least 5% frequency in each country-genepool, we observed that the set of predicted pMHCs differed greatly across countries. Per country, there was a median of 47 (range 1-127) predicted pMHCs, including a median of 6 (range 1-11) HLA alleles and a median of 45.5 (range 1-119) unique peptides (**Table S3**). Still, we identified 20 nine-mer peptides shared by common HLA types across 30 of 77 countries (18 of these peptides correspond to HLA types prevalent in the United States) (**Figure 2**). These peptides spanned 5 genes, including *ORF1ab* (ORF1ab polyprotein, n=14), *S* (Spike glycoprotein, n=2), *M* (membrane protein, n=1), *N* (nucleocapsid protein, n=1), and *ORF3a* (Protein 3b, n=1). Notably, this approach excluded countries in Latin America (such as Brazil and Nicaragua) and in Africa (such as Rwanda and Libya), as HLA types prevalent in these countries do not correspond to this filtered list of peptides.

**Figure 2.**
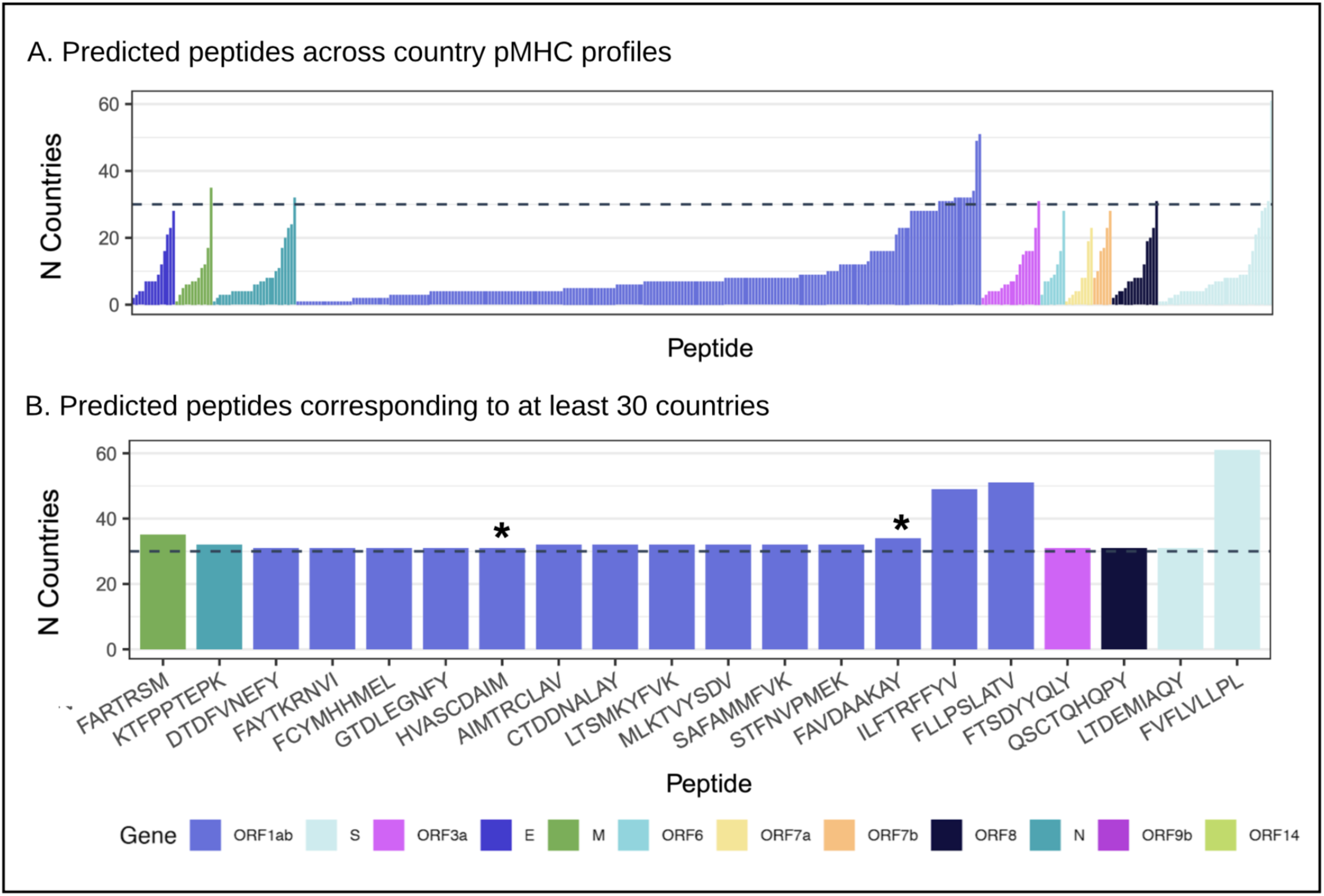
Peptide diversity in country pMHC profiles. Overview of country pMHC profiles, reflecting HLA frequency distributions reported by AFND. Frequency data was filtered to only include alleles with at least 5% frequency for each country. The y-axis indicates the number of country pMHC profiles that included each peptide along the x-axis. Two groups of peptides are shown, according to corresponding SARS-CoV-2 gene: A) peptides that appeared at least once in any country pMHC profile, and B) those that appeared in a minimum of 30 country pMHC profiles. (*) indicates that the peptide was not included in the pMHC profile of the United States.

To improve the global reach of a putative peptide-based vaccine, we utilized a set cover algorithm to determine the smallest set of predicted antigens that covered the maximum number of individuals in each country’s population. An individual was considered “covered” if their simulated class I HLA type was involved in at least one predicted pMHC, and these sets of peptides were denoted as the set cover solutions (SCSs) for the associated population. SCSs were calculated for 77 individual countries and for a “global” population, generated by pooling together the sample populations from all countries, and sampling from this combined pool (n=100,000) without replacement (**Figure S2**, **Table S3**).

Based upon our simulated presentations, SCSs were capable of summarizing predicted pMHCs in at least 90% of the population in 57 countries. Furthermore, in 45 of these 57 countries, SCSs included 30 or fewer peptides (**Figure 3A**). When we evaluated which viral genes were associated with peptides in SCSs, *Orf1ab* contributed the largest number of peptides across all countries. (**Figure 3B**). We filtered peptides to those included in SCSs for at least 30 countries, and identified 19 predicted peptides, spanning 9 genes (**Figure 3C**).

**Figure 3.**
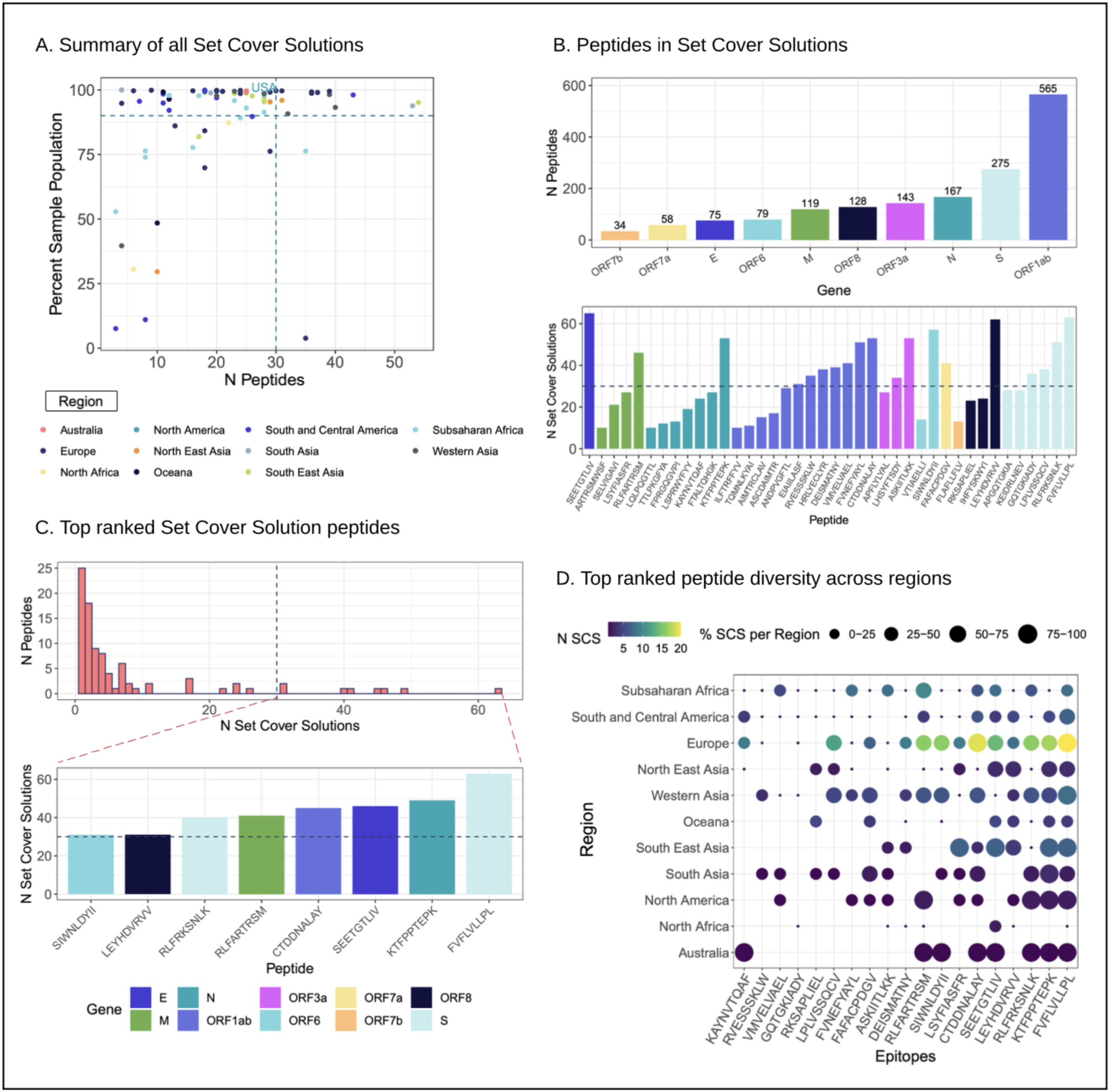
Set Cover Solution peptide summary. A. Summary of SCS results for all 77 countries. The percent of the sample population covered and the number of peptides involved in each SCS is shown, annotated by each country’s corresponding region. The United States is also denoted by text. B. Overview of peptides comprising SCSs. The number of peptides each SARS-Cov-2 protein contributed is shown (top), as well as the number of SCSs individual peptides contributed to (bottom), filtered to show peptides that contributed to a minimum of 10 SCSs. C. Peptides were ranked within each SCS, based upon those associated with the largest cumulative percentage of the population. There were 95 unique peptides comprising the top 10 ranking of all SCSs, and are shown in this figure. The histogram (top panel) shows the number of SCSs associated with each of these peptides. The most recurrent peptides (present in over 30 SCSs) are further shown (bottom panel). D. Geographic distribution of top-ranked peptides, shown in C. Peptides were filtered to those associated with country SCSs spanning at least 5 regions are shown. Each tile is colored by the number of SCSs (i.e. countries) within each global region (y-axis) corresponding to each peptide (x-axis).

The constructed SCSs were also used to prioritize peptides of interest across geographic regions. Peptides were ranked within each SCS, based upon those associated with the largest cumulative percentage of the population. Evaluating the top ten ranked peptides within each SCS (n=95 unique peptides), each was associated with a mean of 7.73 country SCSs (range 1 - 63) (**Figure 3C**). Furthermore, 19 out of 95 top-ranked peptides were associated with SCSs for countries from at least 5 out of 11 global regions (**Figure 4D**). These peptides are of particular interest, as they may be relevant across disparate populations.

**Figure 4.**
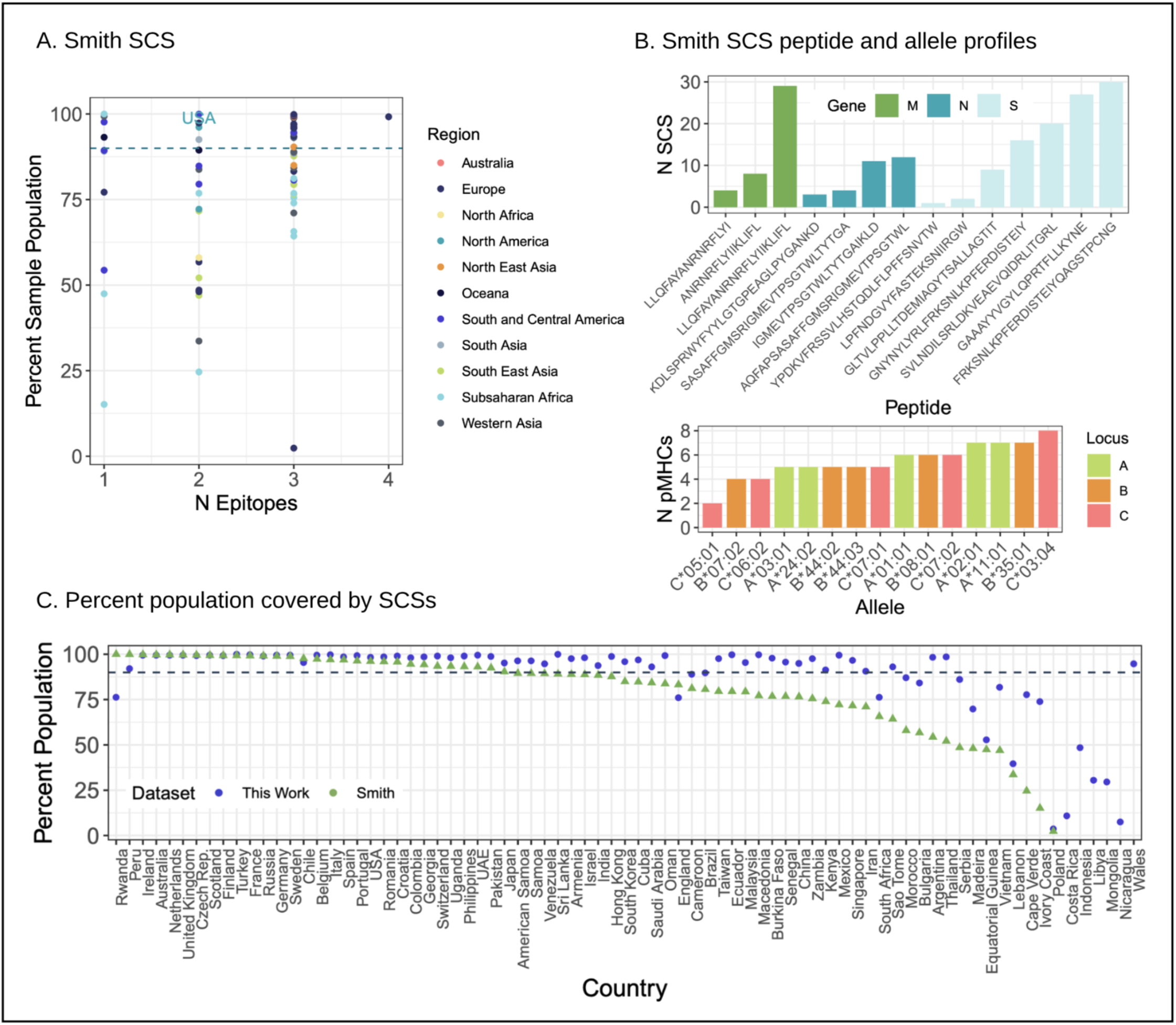
Smith et al. Set Cover Solution summary. A. The Smith et al. dataset was summarized across countries by previously reported epitopes and corresponding HLA types. The number of peptides included and the percent of the sample population covered by each SCS is shown, colored by each country’s corresponding region. The United States is also denoted by text. B. The peptides comprising the Smith SCS are shown (top panel), along with the number of SCSs (y-axis) associated with each peptide and the corresponding SARS-CoV-2 gene (color). The HLA Alleles associated with these peptides is also shown (bottom), according to the number of pMHCs (i.e. peptides) predicted for the corresponding allele. C. The percent of each simulated country (x-axis) population covered by their respective SCSs is shown; SCS from this work is shown in blue, and the SCS results from Smith et al. is shown in green. A 90% cutoff is denoted by the horizontal, dashed line.

We compared the SCSs established from our predicted pMHCs to SCSs generated from a “reference” set of published peptide vaccine candidates ((Grifoni et al., 2020a; Smith et al., 2020; Weiskopf et al., 2020b) based upon highly prevalent HLA types in the United States and overlapping epitopes derived from both CD4 T cell, CD8 T cell, and B cell epitope predictions. The SCSs derived from the reference peptides were relevant in at least 90% of the population across 30 countries, including the United States (**Figure 4A**). Notably, the 14 epitopes that comprised these SCSs were associated with 15 HLA types that were prevalent (at least 5% allelic frequency) across an average of 25 country populations, including HLA-C*03:04, B*35:01, A*11:01, and A*2:01 (**Figure 4B**). In contrast, the SCSs from our dataset were relevant in at least 90% of the population for 57 countries (**Figure 4C**) by including 164 additional predicted peptides associated with 823 additional HLA types (**Table S3**). Thus, the inclusion of these additional HLA types or peptides in development may broaden the global applicability of vaccines.

### Deployment of a user interface to explore the epitome of SARS-CoV-2

To make the predictions generated in this study publicly available and accessible to facilitate experimental validation, we established a user interface to explore the predicted SARS-CoV-2 epitopes in this dataset (https://rstudio-connect.parkerici.org/content/13/). Predicted T cell epitopes can be filtered by features described in this study, including viral gene or protein, peptide length, peptide sequence, HLA gene or specific type, and country (population) HLA allelic frequency. Furthermore, filtered predictions are mapped to other published datasets, including those validated or reported by other groups (Grifoni et al., 2020b; Smith et al., 2020). SCSs generated for this study are also made available through this interface. This tool will serve as a resource for the development of virus specific T cell assays or vaccine design, by considering the global landscape of HLA susceptibility in SARS-CoV-2.

## Discussion

Our study was designed to evaluate the predicted epitope landscape with respect to the SARS-CoV-2 viral proteome across a globally representative set of HLA alleles. We aimed to establish a resource for the scientific community, and have made the entirety of these data publicly available and accessible. This work expands upon recent studies that inferred the epitope landscape of SARS-CoV-2 to either interrogate T cell responses in infected individuals or develop vaccines (Chour et al., 2020; Grifoni et al., 2020a, 2020b; Smith et al., 2020; Weiskopf et al., 2020a). Our pan-HLA approach enabled identification of new HLA contexts for previously proposed and validated peptides, as well as the identification of additional peptides from less prevalent HLA types. Furthermore, the overlap between class I and II predicted pMHCs suggests that some epitopes may be presented to both CD4 and CD8 T cells.

The pan-HLA approach, the inclusion of the entire SARS-CoV-2 proteome, and integration of HLA frequency data from AFND allowed unique evaluation of the regional and global relevance of our predicted pMHC dataset. We establish a set-cover based approach to explore the relevance of our predicted pMHCs across distinct global populations, and use this to construct sets of predicted pMHCs that have putative relevance across 90% of the population in 57 countries. These set cover solutions were superior using our dataset, compared to previously published datasets of peptide-based vaccine candidates, due to the breadth of predicted pMHCs and HLA subtypes.

This dataset and analysis have limitations. Our analysis was restricted to pMHC complexes with predicted binding affinities of less than 500nM. Subsequent analysis did not treat the predicted binding affinities as a continuous variable (i.e. predicted values of 5nM and 400nM were treated similarly in the remaining analysis). In the absence of experimental validation, we did not try to over delineate the association between HLA diversity and the predicted binding affinity. Furthermore, utilizing a threshold of 500nM may result in underestimating the number of alleles associated with the predicted antigenic peptides. Our predictions were limited to 9-mers and 15-mers, which represent most but not all reported HLA class I and class II binding peptides. Our data also does not account for either the quantity or timing of viral protein expression in a host cell, both of which can impact the immunogenicity of predicted epitopes (Croft et al., 2019). Finally, analysis of global population frequencies was restricted to a limited number of HLA alleles and countries. While AFND is the most comprehensive database summarizing the population frequencies of HLA haplotypes, it is far from complete. Frequencies are reported for 73, 73, and 49 countries for genes *HLA-A, -B,* and *-C*, respectively. In addition, the number of alleles reported for each gene is variable across countries, ranging from 1-1,498.

In summary, our resource provides a pan-HLA tool for those seeking to study SARS-CoV-2 or vaccine-induced T cell responses. In addition, our strategy enables the identification of sets of class I peptides either within or across countries, an important consideration for vaccine design. For these reasons we have made our calculations available in full and have also developed a user-friendly web-app to enable exploration of these data at https://rstudio-connect.parkerici.org/content/13/. SARS-CoV-2 is an ongoing pandemic and these resources will be updated as further peptide validation becomes available.

## Author Contributions

K.M.C., G.S., and A.K. conceived experiments. K.M.C. and A.K. supervised the study. K.M.C., G.S., D.K.W., A.R., and A.K. designed the experiments. K.M.C. and G.S. performed data processing and analysis. K.M.C., G.S., and A.K. wrote the manuscript, and all authors contributed to final revisions of the manuscript.

## Acknowledgments

We are grateful to John Wherry (University of Pennsylvania, Philadelphia, PA) and Bonaventura Clotet, Julia Garcia Prado and Christian Brander (IrsiCaixa Foundation, Barcelona, Spain) for valuable feedback on the manuscript. The computational resources for this study were provided by the Parker Institute for Cancer Immunotherapy (PICI). K.M.C. is supported by the UCLA Tumor Immunology Training Grant (NIH T32CA009120) and the Cancer Research Institute (CRI) Irvington Postdoctoral Fellowship Program. A.K. is supported by the UCLA CTSI KL2 Award (NCATS TR001882) and Sarcoma Alliance for Research Through Collaboration Career Enhancement Program. A.R. is supported by R35 CA197633 and The Ressler Family Fund, and is a member researcher at PICI.

## STAR METHODS

### Resource Availability

#### Lead Contact

Further information and requests for resources and reagents should be directed to and will be fulfilled by the Lead Contact, Katie Campbell.

#### Materials Availability

##### Data and Code Availability

The results of this study are available in a public Google bucket through the following link: https://console.cloud.google.com/storage/browser/pici-covid19-data-resources (gs://pici-covid19-data-resources). This bucket contains all of the unfiltered peptide binding predictions and the Supplemental Tables corresponding to this document. The filtered peptide binding predictions and set cover solutions can be explored using the interactive Shiny app at https://rstudio-connect.parkerici.org/content/13/. All filtered data can be exported from this web interface. The code for this manuscript and the Shiny App is available in the public github repository https://github.com/kcampbel/neocovid-app.

### Method Details

#### Data acquisition

The NCBI Virus resource (https://www.ncbi.nlm.nih.gov/labs/virus/vssi/) was used to obtain all annotated protein sequences for SARS-CoV-2 on March 15, 2020. This dataset spanned 166 genotypes, but protein sequences were summarized by the corresponding UniProt annotation for the SARS-CoV-2 proteome (UP000464024). All possible 9-mer and 15-mer peptides were obtained from the entire viral proteome for Class I and Class II peptide binding predictions, respectively.

#### Peptide-MHC binding predictions

Peptide binding predictions were performed using the pVACbind tool from pVACtools and executed using the griffithlab/pvactools:1.5.7 (https://hub.docker.com/r/griffithlab/pvactools/) Docker image. Class I prediction algorithms included MHCflurry (v1.6.0) (O’Donnell et al., 2018), MHCnuggets (v2.3) (Shao et al., 2020), NetMHC (v4.0) (Andreatta and Nielsen, 2016), PickPocket (v1.1) (Zhang et al., 2009), SMM (v1.0) (Peters and Sette, 2005), and SMMPMBEC (v1.0) (Kim et al., 2009). Class II prediction algorithms included NetMHCIIpan (v4.0) (Reynisson et al., 2020), SMMalign (v1.1) (Nielsen et al., 2007), NNalign (v2.3) (Nielsen and Andreatta, 2017), and MHCnuggets (v2.3) (Shao et al., 2020).

HLA alleles were chosen by running the command `pvacseq valid_alleles` and filtering out any non-expressed or null HLA alleles (those ending with the “N” suffix), resulting in 9,360 Class I HLA proteins and 8,445 Class II HLA alleles. It is important to note that for Class II predictions, some algorithms include inputs of either individual HLA alleles (e.g. DPB1*01:01) or combinations of HLA alleles (e.g. DPA1*01:03-DPB1*01:01), since two HLA proteins pair together for Class II antigen presentation. All available individual (n=3,484) or combinations of (n=4,961) Class II HLA alleles were used for input.

Class I predictions were performed using the following command:

$ /opt/conda/bin/pvacbind/run ${fasta} ${hla} ${hla} MHCflurry MHCnuggetsI NetMHC PickPocket SMM SMMPMBEC tmp/ -e 9 --iedb-install-directory /opt/iedb --net-chop-method cterm --netmhc-stab

Class II predictions were performed using the following command:

$ /opt/conda/bin/pvacbind/run ${fasta} ${hla} ${hla} NetMHCIIpan SMMalign NNalign MHCnuggetsII tmp/ - e 15 --iedb-install-directory /opt/iedb --net-chop-method cterm --netmhc-stab

Where ${fasta} was the protein sequence fasta containing the SARS-CoV-2 proteome, obtained at NCBI Virus. The pVACbind tool was performed individually across the union of HLA alleles available for all algorithms, and each allele was specified by the ${hla} input in the command. The filtered results, containing peptide-MHC complexes with predicted Median Score (nM) less than 500nM, were aggregated for the final dataset.

#### Population frequencies of HLA types

### Country Populations

Population frequencies of HLA alleles were obtained from the Allele Frequency Net Database (Gonzalez-Galarza et al., 2020). The database contains HLA Frequency data for Class I and Class II alleles across 1,028 distinct populations. Populations whose net frequency data exceeded 1 at a given allele were excluded from this analysis, as well as populations that did not report frequencies for alleles with 2 or 3 fields. Because these populations are highly granular (i.e. “USA San Francisco Caucasian”), we aggregated them into 98 populations by country; 77 had data for Class I alleles. This was done by (1) assigning each population to a country using the first word from each population name, (2) calculating the country “sample size” by summing the sample sizes of distinct populations, and (3) calculating HLA frequencies within each country population using **Formula 1**.

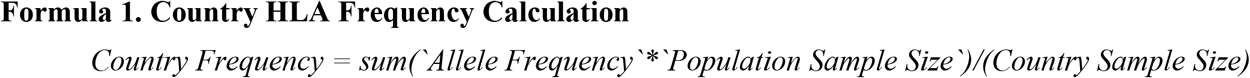

Sample populations were generated from this country-frequency data for 77 countries. This was done by sampling 2n alleles from each country gene pool for each Class I allele with reported data, where n represents the number of simulated individuals in the sample population (n = 100,000). The probability of selecting an allele for each sample population is equal to the frequency of that allele reported for each given country. The simulated populations used for analysis are available in the github repository (https://github.com/kcampbel/neocovid-app). These populations of simulated genotypes were then merged with our Class I predictions to create a pMHC profile for each country, based on each country’s reported allele frequencies.

### Global Population

A simulated “global population” was generated by first aggregating all 77 country sample populations, and then sampling from this pool 2n times to create a sample population of n individuals (n = 100,000). This ensured that each country would be represented in this global population with equal probability regardless of sample size, such that the global population was not further biased towards the United States and European countries. It should be noted that consequently, this global population does not reflect true global HLA frequencies, which would require consideration of true country size.

#### Set Cover Solutions

Given a universal set of n elements (U), a collection of subsets of U (S), and the associated cost of each subset in S, the set cover problem is to identify I, the minimal subcollection of S, whose union equates to U and minimizes the total cost (Karp, 1972). The greedy algorithm addresses this problem by iteratively adding elements of U to I until all subsets in S are covered (Vazirani, 2013). This problem is NP-hard, so a logN approximate solution was used.

That is, for a simulated population X,

U = all individuals in X covered by at least one pMHC
S_i = all individuals in X covered by the pMHC with Epitope i
S = set of S_i whose union spans all individuals in U
cost(S_i) = 1 for all Epitopes i

The solution (I) represents the smallest set of epitopes whose union covers the largest portion of population X.

### Quantification and Statistical Analysis

Data analysis and visualization was performed in R using the tidyverse packages [REF]. The neoCOVID Explorer application was developed using the Shiny R package [REF] and deployed using RStudio-Connect.

### Key Resources Table

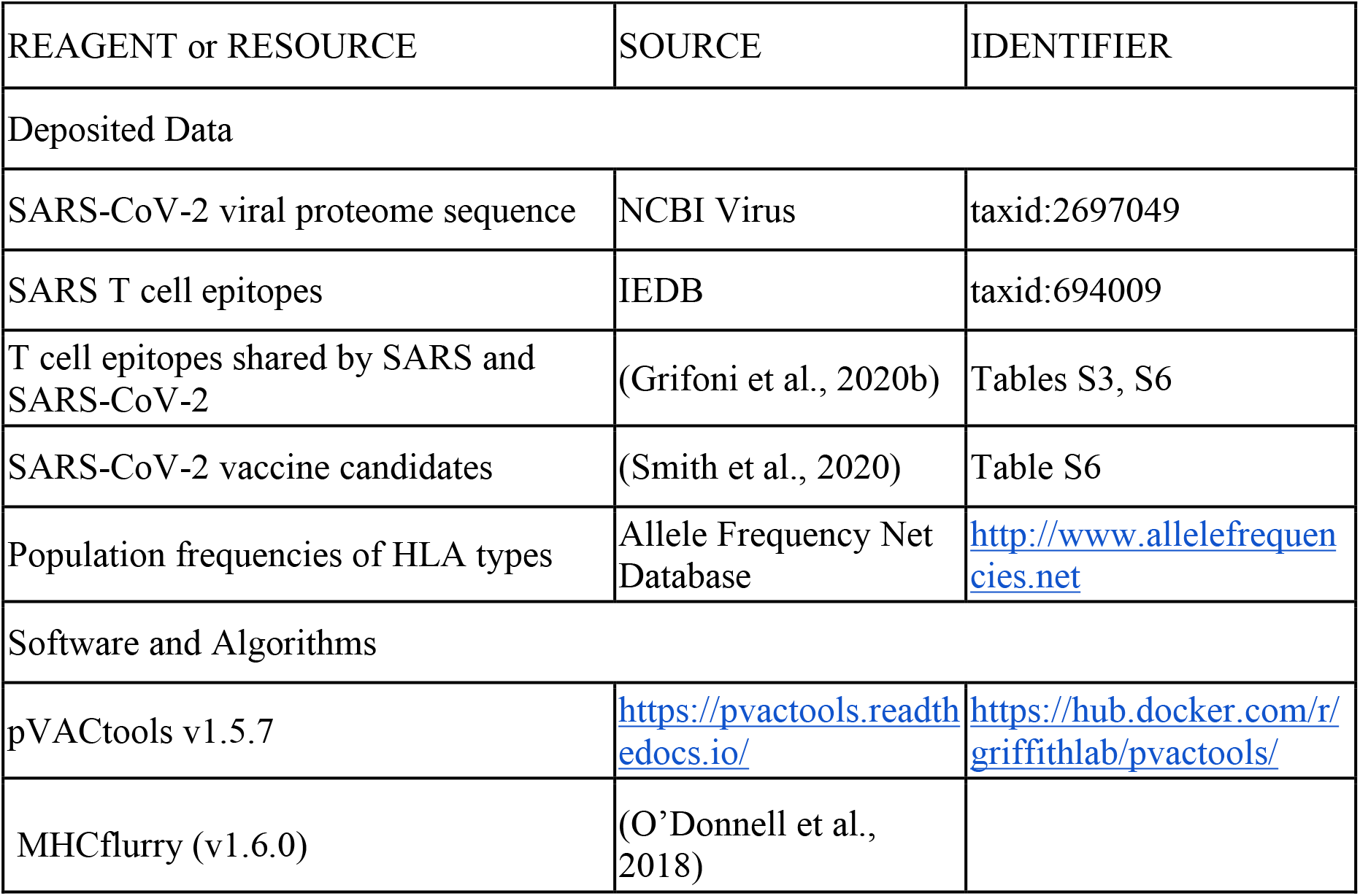

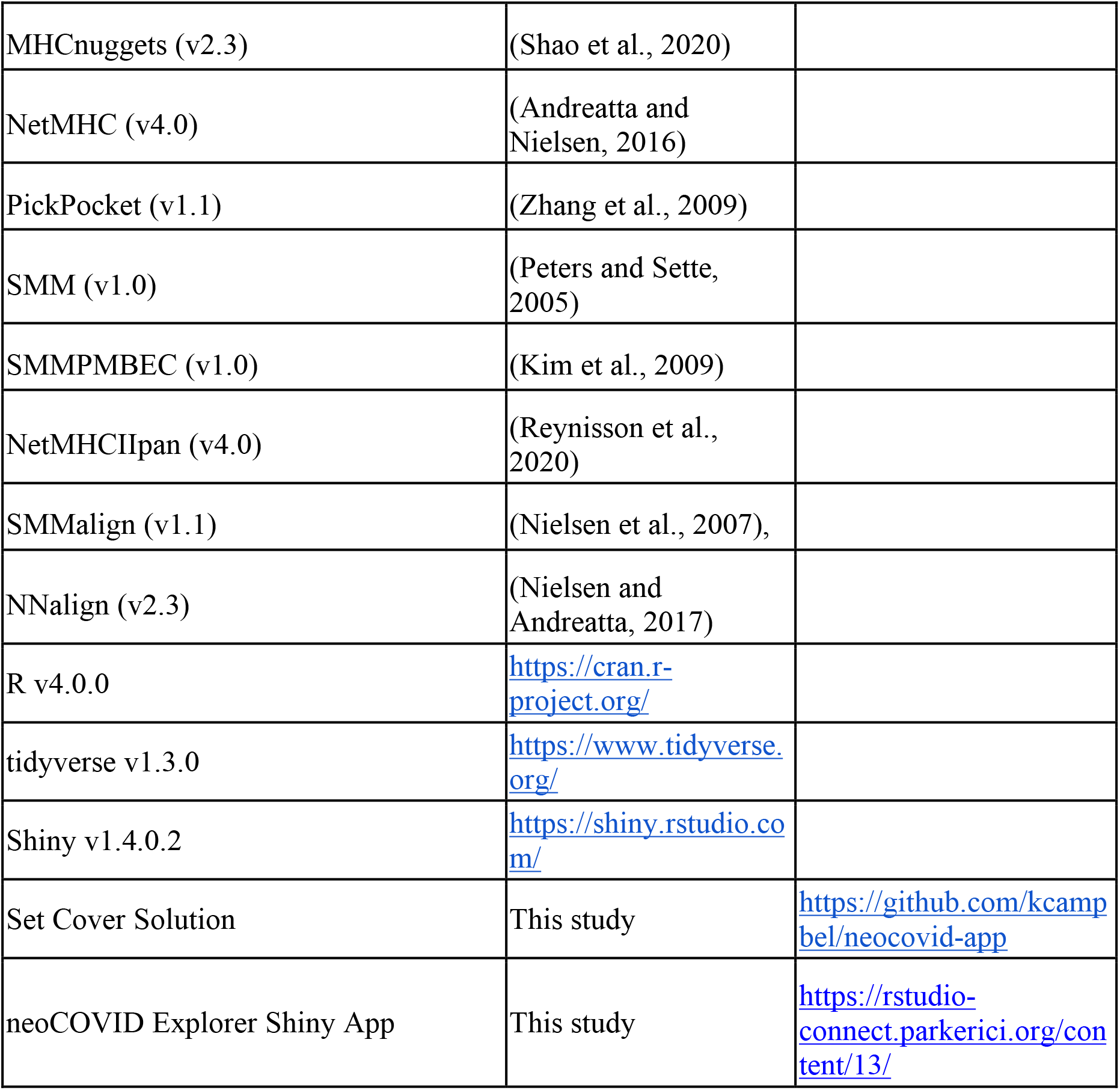

