## Supplementary Appendix for "Prioritization of SARS-CoV-2 epitopes using a pan-HLA and global population inference approach"

**SUPPLEMENTAL INFORMATION**

*SUPPLEMENTAL FIGURES*

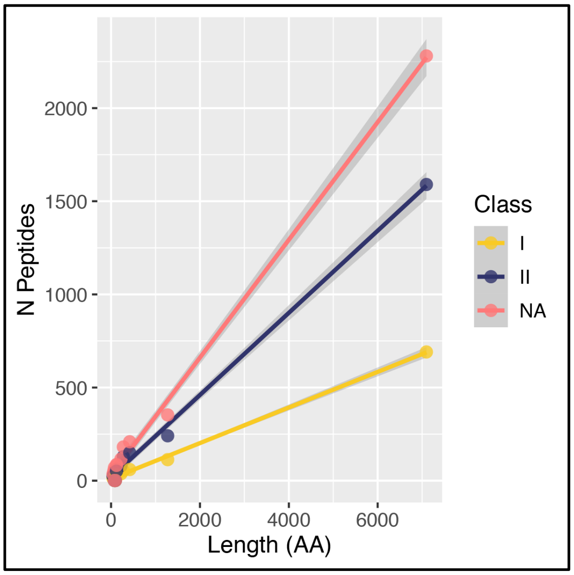


**Figure S1. Correlation of predicted number of peptides with gene length**

The number of unique peptides (y-axis), corresponding to pMHCs with predicted binding affinity of <500nM, was compared to the length, in number of amino acids, (x-axis) of each viral gene. Results are colored by either Class I, II, or Total (sum of Class I/II) peptides.

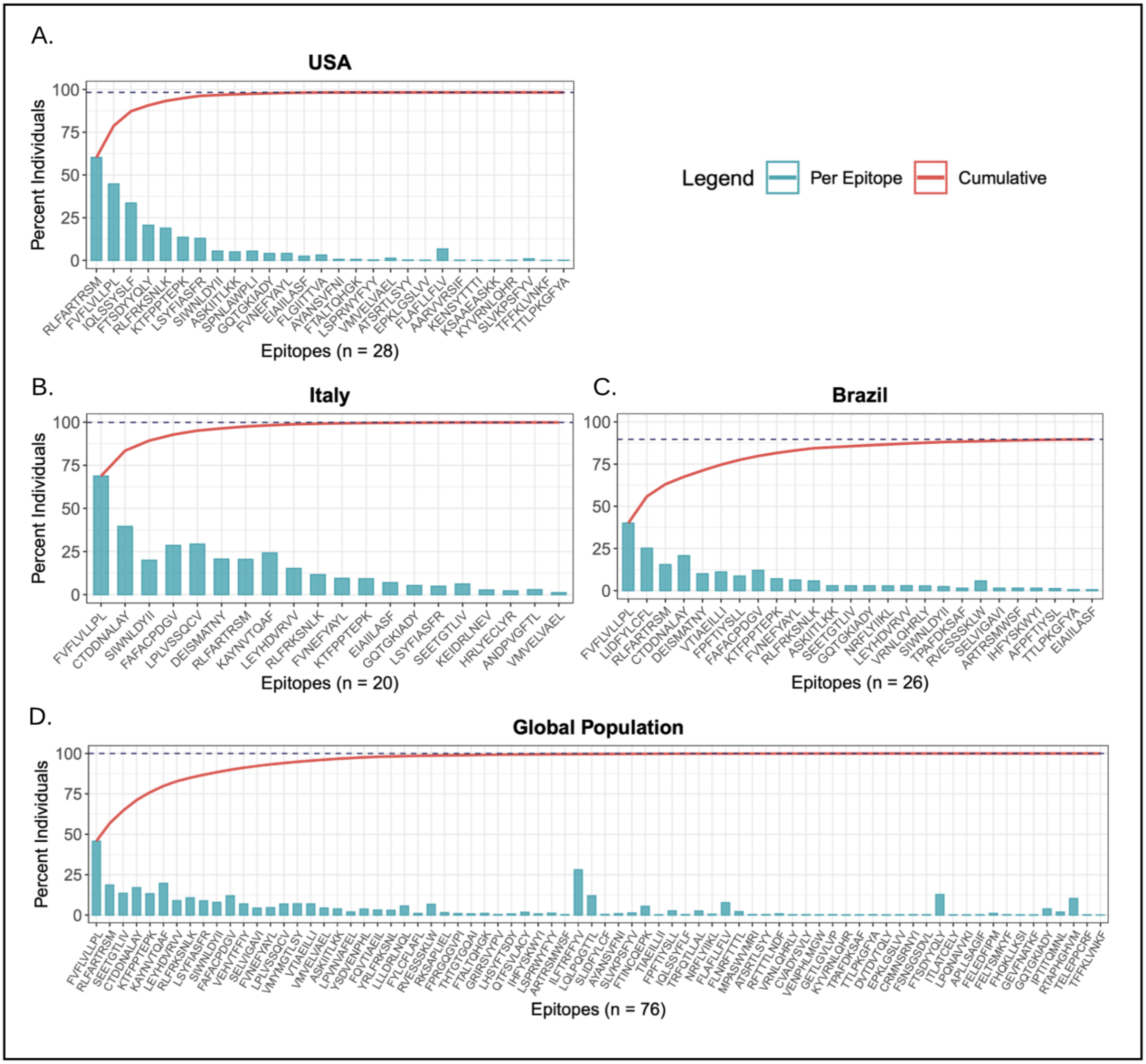


**Figure S2. Set Cover Solution results**

SCS results are shown for the simulated A) USA, B) Italy, C) Brazil, and D) Global (right) populations. The percent of the simulated population covered by each peptide alone is shown in blue (bar), and the percent covered by the cumulative sum of peptides (left to right) is shown in red (line). Peptides appear along the x-axis according to the sequence in which they were returned by the algorithmic solution (denoted as their SCS "rank"). For the specific peptide sequence for all SCS, see Table S6.

*SUPPLEMENTAL TABLES*

**Table S1. Filtered peptide binding predictions**

This table includes peptide-MHC complexes with predicted binding affinities of <500nM. The columns 'IEDB' and 'Grifoni' list the peptides either overlapping or nested within either of these datasets.

**Table S2. HLA Frequency Data**

This table includes HLA Frequency data aggregated across countries from AFND.

**Table S3. Set Cover Solutions**

This table includes the Set Cover Solutions generated for all countries with available AFND data.
